# GLS1 is a protective factor rather than a molecular target in ARID1A-mutated ovarian clear cell carcinoma

**DOI:** 10.1101/2021.09.03.457161

**Authors:** Valentino Clemente, Andrew Nelson, Britt Erickson, Ruth Baker, Nathan Rubin, Mahmoud Khalifa, Asumi Hoshino, Mihir Shetty, Emil Lou, Martina Bazzaro

## Abstract

Targeting glutamine metabolism has emerged as a novel therapeutic strategy for several human cancers, including ovarian cancer. The primary target of this approach is the kidney isoform of glutaminase, glutaminase 1 (GLS1), a key enzyme in glutamine metabolism that is overexpressed in several human cancers. A first-in-class inhibitor of GLS1, called CB839 (Telaglenastat), has been investigated in several clinical trials, with promising results. The first clinical trial of CB839 in platinum-resistant ovarian cancer patients is forthcoming. *ARID1A*-mutated ovarian clear cell carcinoma (OCCC) is a relatively indolent and chemoresistant ovarian cancer histotype. In OCCC-derived cells in vitro and mouse models, loss of ARID1A leads to upregulation of GLS1. Thus, targeting of GLS1 with CB839 has been suggested as a targeted approach for OCCC patients with tumors harboring *ARID1A*-mutations. Here, we investigated whether GLS1 is differentially expressed between OCCC patients whose tumors are ARID1A positive and patients whose tumors are ARID1A negative. In clinical specimens of OCCC, we found that GLS1 overexpression was not correlated with ARID1A loss. In addition, GLS1 overexpression was associated with better clinical outcomes. Our findings suggest that GLS1 expression in OCCC may be a protective factor and that caution should be taken when considering the use of CB839 to treat OCCC patients.

## Introduction

Ovarian clear cell carcinoma (OCCC) is an indolent form of ovarian cancer associated with a poor prognosis. The principal reason for such a dismal prognosis is resistance to standard-of-care chemotherapy used in general for ovarian carcinomas, including taxol- and platinum-based agents^1^. OCCC is characterized by a specific subset of genetic mutations, the most frequent one being inactivating mutations (protein loss) of *ARID1A*, which is found in approximately 50% of patients^1–5^. ARID1A is a member of the SWI/SNF (SWIft/Sucrose Non-Fermentable) complex of chromatin remodelers and is considered a tumor suppressor^5^. Mutations in genes coding for members of the SWI/SNF complex are found in approximately 20% of all human cancers^6^. OCCC patients carrying *ARID1A* mutations, including patients that are diagnosed at early stages, have worse prognoses than patients without *ARID1A* mutations^5,7–15^. Hence, there is a notable urgency to discover new and more effective treatments based on the molecular dependencies of *ARID1A*-mutated OCCC.

We and others have contributed to the understanding of how the SWI/SNF remodelling complex controls the energetic metabolism of mammalian cells, including the mitochondrial metabolism of cancer cells. In fact, we have recently shown that ARID1A loss is associated with higher dependency upon mitochondrial respiration and selective sensitivity to its inhibition, both *in vitro* and in a preclinical model of *ARID1A*-mutated OCCC^16^. This result is consistent with a previous study showing that OCCC-derived cells have higher mitochondrial respiration as compared to ovarian cancer cell lines derived from other histotypes.^17^ It is also consistent with another study showing that ARID1A loss leads to selective sensitivity to ROS-inducing agents in OCCC cells^18^. Furthermore, a recent report shows that, in ARID1A knock-out cells, increased glutamine metabolism and GLS1, the key regulating enzyme in glutamine metabolism, are responsible for fueling mitochondrial respiration^19^. Thus, inhibition of GLS1 with the novel inhibitor CB839 (Telaglenastat) has been proposed as a novel strategy for the treatment of *ARID1A*-mutated OCCC^19^. This is conceptually supported by findings from our team^16^ and others^20–24^ showing that in OCCC, ARID1A loss is followed by increased expression levels of c-Myc, which is known to be an important regulator of GLS1 expression.^25–47^. In this scenario and given that the mitochondrial pathways involve a number or proteins organized in complexes and super complexes, GLS1 would represent an ideal marker and molecular target for *ARID1A*-mutated tumors as it represents the “bottleneck” for mitochondrial glutamine metabolism. Hence, there is strong enthusiasm and scientific rationale for the launch of a forthcoming clinical trial utilizing CB839 in ovarian cancer including OCCC^48^.

In this study, we addressed the question of whether GLS1 is differentially expressed between OCCC patients whose tumors are ARID1A positive and patients whose tumors are ARID1A negative. This question is relevant because it would provide a biomarker for stratification of OCCC patients based on the likelihood to respond to CB839 treatment. We found that in clinical specimens of OCCC GLS1 overexpression was not correlated to ARID1A loss and, on the contrary, was associated with better clinical outcome. This result suggests that in OCCC GLS1 may be a protective factor and that caution should be taken when considering the use of CB839 to treat OCCC patients.

## Materials and methods

### Antibodies

Anti-ARID1A (HPA005456, Sigma-Aldrich, St. Louis, MO, USA) and anti-GLS1 (ab156876, clone EP7212, Abcam, Cambridge, MA, USA).

### Human Subjects

Archival tissues were used with approval from the Institutional Review Board (IRB) of the University of Minnesota (STUDY00006529). Demographic information and patient characteristics are reported in Table 1.

### Immunohistochemistry

Representative cores from the formalin fixed paraffin embedded blocks of archival tissues were selected and arranged in tissue micro-arrays (TMAs). Five-micron thick, formalin-fixed, paraffin-embedded TMA sections were deparaffinized and rehydrated by sequential washing with xylene, 100% ethanol, 95% ethanol, 80% ethanol, and PBS. Antigen retrieval was then carried out with 1X Reveal Decloaker (Biocare Medical, Pacheco, CA, USA) in a vegetable steamer for 30 min at 100 °C, before blocking the slides with Background Sniper (BS966H, Biocare Medical, Pacheco, CA, USA) for 13 min at room temperature. After washing with PBS, sections were incubated with anti-ARID1A (1:250 dilution) or ant-GLS1 (1:100 dilution) antibodies overnight at 4 °C. After washing twice with PBS, the sections were incubated with Biotin-SP-conjugated AffiniPure Goat Anti-Rabbit IgG (111-065-003, Jackson ImmunoResearch Laboratories, West Grove, PA, USA) at a dilution of 1:200 for 30 min at room temperature followed by incubation with horseradish peroxidase streptavidin at a dilution of 1:125 (405210, BioLegend, San Diego, CA, USA) for 30 min at room temperature. After the staining was developed with 3,3′-diaminobenzidine (926506, BioLegend, San Diego, CA, USA) for 3 min, slides were counterstained with Harris’ hematoxylin. Immunostained slides were reviewed by a panel of two investigators blinded to the clinical outcome of the corresponding patients. ARID1A immunoreactivity was scored using an immunoreactive score (IRS), and a cutoff of <5.5 was considered predictive of mutation, as previously described^49^. GLS1 immunoreactivity was scored using an H-score: staining intensities (0=no staining, 1=weak, yellow staining, 2=yellow/brown, 3=brown) were multiplied by their respective percentages of cells stained (final range: 1 to 300). Brightfield images were acquired with a Zeiss Axio Scan.Z1 system (Zeiss, Thornwood, NY, USA) at 40× magnification.

### Statistical analysis

Patient demographic and clinical measures were summarized and compared between ARID1A and GLS1 categorical groups using either one-way ANOVA or 2 sample *t*-tests for continuous measures and Fisher’s Exact tests for categorical measures. In this analysis ARID1A was treated as a binary variable using the previously mentioned cut-off of 5.5, while GLS1 categorical groups were created based on low (<100), moderate (<200, ≥100) and high (≥200) expression levels. Survival curves were generated using the Kaplan-Meier method. Univariate Cox regression models were used to determine the statistical significance of ARID1A and GLS1 on progression-free survival (PFS), cancer-specific survival (CSS), and overall survival (OS). After identifying possible confounders, models were also generated to adjust for factors such as stage and endometriosis^50^. In the survival analyses, GLS1 was treated as a continuous variable. R (Version 3.4.1, The R Foundation for Statistical Computing) was used for demographic and survival analyses, while graphs were obtained using GraphPad Prism (version 8.4.3, San Diego, CA, USA). ARID1A status (< or >5.5) and GLS1 H-score distributions were analyzed using GraphPad Prism. P-values of <0.05 were considered statistically significant.

## Results

### ARID1A loss negatively correlates with the prognosis of OCCC patients

Studies conducted in both OCCC and in endometrial carcinoma have shown that *ARID1A* mutations are found in 50 and 30% of patients, respectively^2,5,9,15,49,51^ and that patients with tumors carrying *ARID1A* mutations have worse prognosis as compared to patients who do not. This finding is true even for patients diagnosed with early-stage disease^7–15^. Thus, we first validated our OCCC clinical cohort by performing ARID1A immunohistochemical staining (IHC) (Figure 1a) to correlate ARID1A expression levels with clinical outcome. Patient demographics and characteristics are listed in Table 1. Importantly, IHC is the most commonly used technique to evaluate ARID1A expression levels^5^. This method has been shown to reliably predict *ARID1A* mutations in 76% of cases when using a no immunoreactivity cut-off^2^ and with 100% sensitivity and specificity when using an IRS <5.5 cut-off^49^, which is what we used in this study. As shown in Figure 1b, we found a distribution of the scores similar to what has been previously shown^49^ with 19 ARID1A-negative samples out of 58 (34.5%). This result is consistent with the prevalence of *ARID1A* mutations, which is normally in the range between 30-70% of OCCC cases^2,5,15,49,51^. Notably, we found that loss of ARID1A occurred more frequently in patients with endometriosis found during pathological diagnosis, and also had a tendency to present at lower stages (Table 1), which is consistent with the fact that ARID1A mutations are frequently associated with endometriosis^1,2,52–57^.

**Figure 1.**
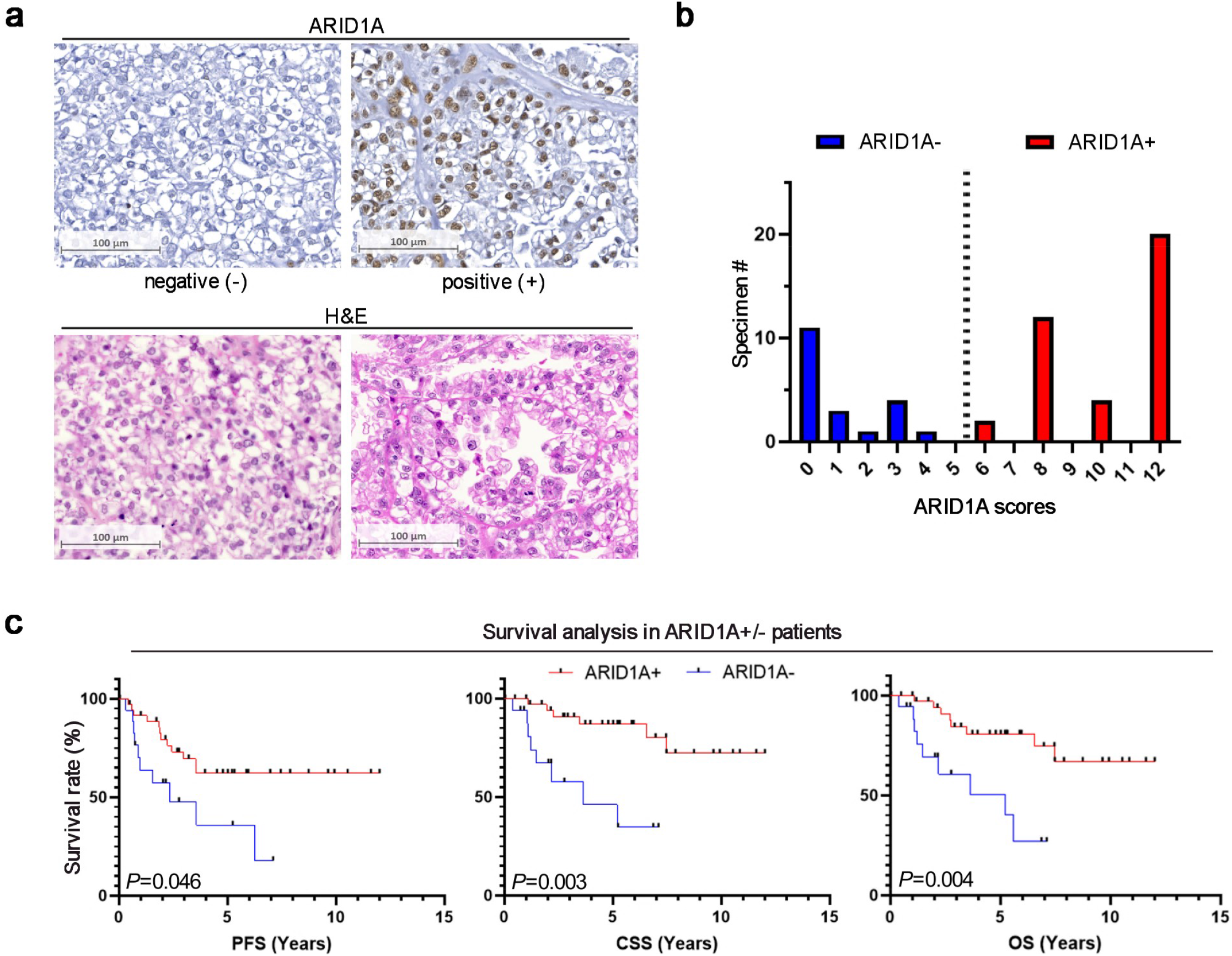
ARID1A loss negatively correlates with the prognosis of OCCC patients. **a**. *Top*, representative images of ARID1A negative or positive OCCC clinical specimens and their respective H&E staining (*bottom*). **b**. Frequency distribution of the immunoreactivity scores for ARID1A; the dotted line represents the cut-off used to discriminate between ARID1A positive (+) and ARID1A negative (-) patients. **c**. Survival curves of ARID1A+ *vs* ARID1A-patients. From the left to the right: progression free survival, PFS (36 vs 19; *P=*0.046; hazard ratio, HR=0.43; 95% confidence interval, CI, 0.19 to 0.98), cancer specific survival, CSS (36 vs 18; *P=*0.003; HR=0.18; 95% CI 0.06 to 0.55), overall survival, OS (36 vs 19; *P=*0.004; HR=0.23, 95% CI 0.09, 0.63).

We then sought to determine the correlation between ARID1A expression and clinical outcome. As shown in Figure 1c, we found a significant and strong detrimental effect for ARID1A loss on PFS (*P*=0.046), CSS (*P*=0.003), and OS (*P*=0.004). Importantly, the correlation between ARID1A loss and worse outcome remained significant even after adjusting for stage and endometriosis in multivariate analysis (Table 2) (PFS, *P*=0.004; CSS, *P*<0.001; OS, *P*<0.001). In fact, consistent with other studies^9^ an even larger effect of ARID1A loss on survival was found in stage I/II patients (Figure 2a) (PFS, *P*=0.034; CSS, *P*<0.001; OS, *P*=0.009), with early stage ARID1A-negative patients having a prognosis similar to that of III/IV ARID1A-positive patients (Figure 2b) (PFS, *P*=0.235; CSS, *P*=0.994; OS, *P*=0.899). Taken together, these results demonstrate that our cohort can reliably be used to investigate the correlation between ARID1A status and GLS1.

**Figure 2.**
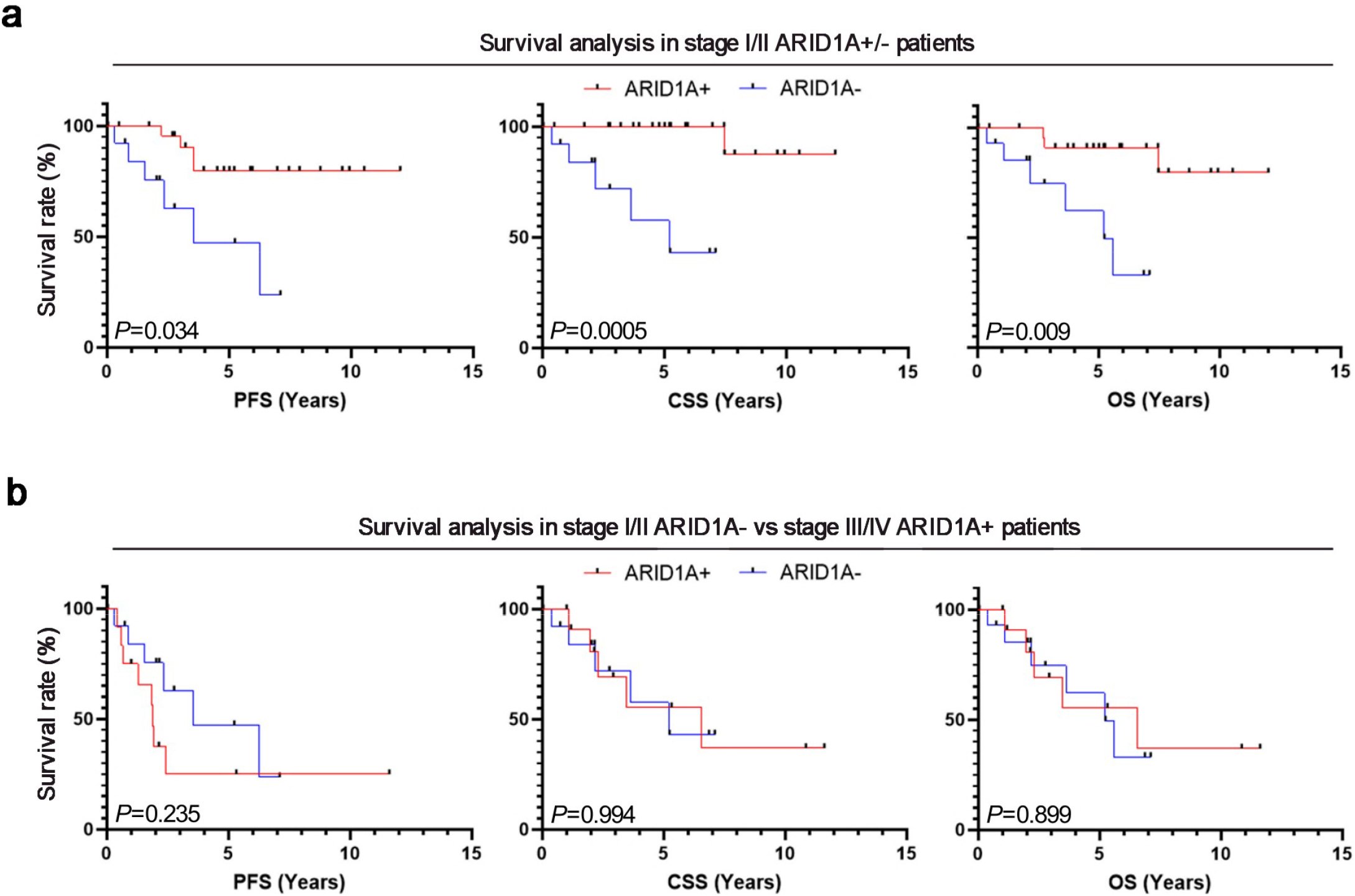
Stage I/II ARID1A negative patients have the same prognoses as stage III/IV ARID1A positive patients. **a**. Survival curves of ARID1A+ (red) vs ARID1A-(blue) stage I/II patients. From the left to the right: PFS (24 vs 15; *P=*0.034; HR=0.29, 95% CI 0.09 to 0.91), CSS (24 vs 14; *P=*0,0005; HR=17,27; 95% CI, 2,538 to 117,6 – Mantel-Cox’s logrank test), OS (24 vs 15; *P=*0.009; HR=0.12, 95% CI 0.02 to 0.59). **b**. Survival curves of stage III/IV ARID1A+ (red) vs stage I/II ARID1A-(blue) patients. From the left to the right: PFS (12 vs 15; *P=*0.235; HR 0.52, 95 CI 0.18 to 1.52), CSS (12 vs 14; *P=*0.994; HR= 1.01, 95% CI 0.29 to 3.49), OS (12 vs 15; *P=*0.899; HR=1.08, 95% CI 0.33 to 3.56).

### GLS1 expression is negatively correlated with ARID1A loss in OCCC

Recent reports have shown that GLS1 is overexpressed in both OCCC and high grade-serous carcinoma (HGSC) of the ovaries, compared to cells derived from normal surface epithelium of the ovaries and fallopian tubes^25^. Furthermore, in HGSC, GLS1 is overexpressed in clinical tumor specimens from patients who are chemoresistant versus patients whose tumors are responsive to chemotherapy^25^. Thus, GLS1 has been proposed as a rational molecular target for chemoresistant ovarian cancer. Here, we sought to assess whether GLS1 is differentially expressed between OCCC patients whose tumors are ARID1A positive and patients whose tumors are ARID1A negative. We found that GLS1 is not overexpressed in clinical specimens of OCCC that are negative for ARID1A (Figure 3a). We found that ARID1A positive tumors have higher levels of GLS1 as compared to patients who are ARID1A negative (Figure 3b and c) (*P*=0.001). This finding is novel and unanticipated. We then compared GLS1 expression levels between ARID1A-negative OCCC and normal surface epithelia from ovaries and fallopian tubes. We found that GLS1 expression in normal epithelia is similar to expression levels detected in ARID1A-negative OCCC (Figure 4 and Supplementary Figure 1a) (*P*=0.828). Furthermore, GLS1 levels were not correlated with other clinicopathological parameters, suggesting that the ARID1A status was the only determinant of GLS1 levels in our cohort (Table 3).

**Figure 3.**
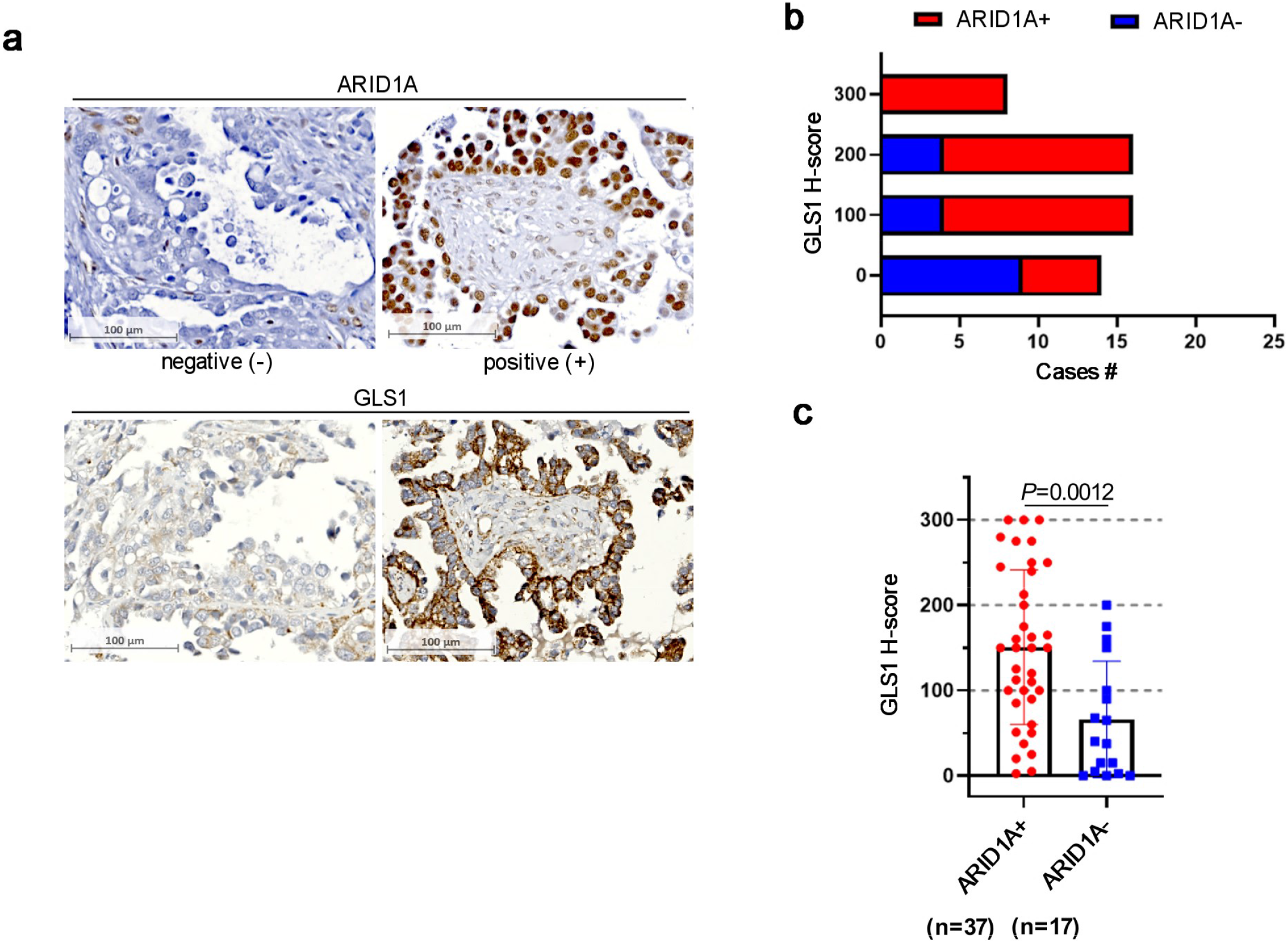
GLS1 expression is negatively correlated with ARID1A loss in OCCC. **a**. *Top*, representative images of GLS1 immunohistochemical staining in ARID1A positive and ARID1A negative OCCC specimens; *bottom*, respective ARID1A immunostainings. **b**. Frequency distribution of the GLS1 H-scores in ARID1A+ and ARID1A-OCCC. **c**. Expression levels of GLS1 in ARID1A+ and ARID1A-OCCC (ARID1A+ *vs* ARID1A-: *P=*0.0012). n= number of clinical specimens per group

**Figure 4.**
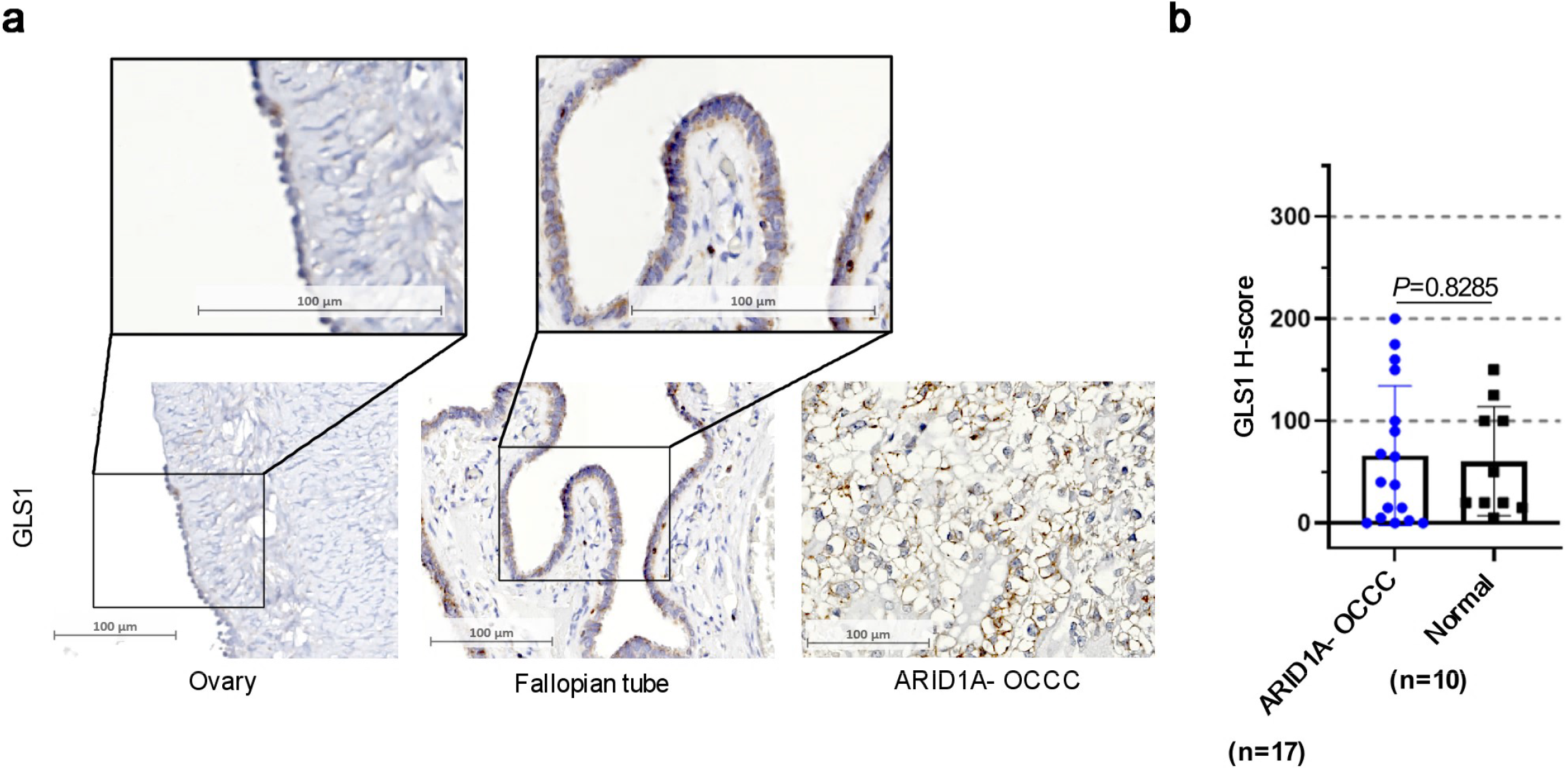
GLS1 expression in ARID1A negative OCCC is similar to the one of normal tissues. **a**. *Bottom left and its inset*, representative image of GLS1 staining in normal ovary. *Bottom center and its inset*, representative image of GLS1 staining in normal Fallopian tube. *Bottom right*, representative image of GLS1 staining in ARID1A negative (-) OCCC. **b**. Expression levels of GLS1 in ARID1A negative OCCC and in normal tissues. (ARID1A-*vs* normal: *P=*0.8285). n=number of clinical specimens per group.

### GLS1 overexpression may be a protective factor in OCCC

We next sought to determine the prognostic significance of GLS1 expression levels in our cohort of OCCC clinical specimens. We found that, while the GLS1 expression levels were not correlated to any specific demographic characteristic of the patients, high GLS1 levels were associated with better survival (Figure 5a) (PFS, *P*=0.013; CSS, *P*=0.014; OS, *P*=0.021). Next, because we found that loss of ARID1A immunoreactivity acted as a negative prognostic factor in our set of patients, and that this correlated with low levels of GLS1, we included the ARID1A status in multivariate analysis to exclude the hypothesis that it may have acted as a confounder. Both the factors tended to remain significant, but ARID1A did not reach significance for PFS (PFS, *P*=0.271; CSS, *P*=0.026; OS, *P*=0.042), and GLS1 only showed a trend toward improved OS (PFS, *P*=0.031; CSS, *P*=0.033; OS, *P*=0.064) (Table 4). Furthermore, when analyzing the effect of GLS1 on the prognosis of the ARID1A+ and ARID1A-subpopulations individually, all the survival curves showed a similar trend to the general population (Figure 4b and c) (ARID1A+: PFS, *P*=0.023; CSS, *P*=0.036; OS, *P*=0.093), but significance was lost in the ARID1A-subgroup (Figure 4c) (PFS, *P*=0.487; CSS, *P*=0.647; OS, *P*=0.511), probably due to the very small size of the moderate and high expression groups (n=4 and 1, respectively).

**Figure 5.**
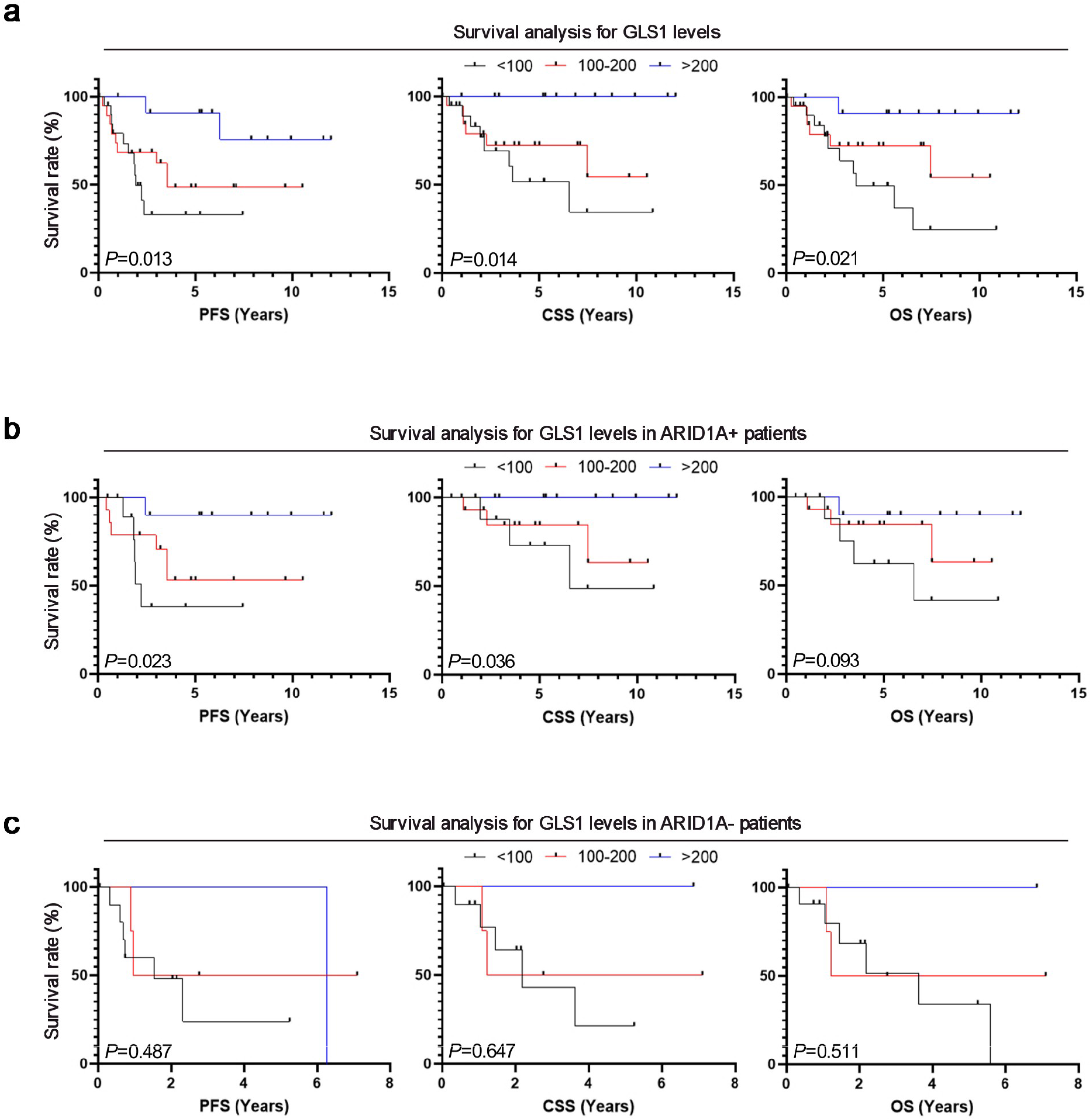
GLS1 overexpression may be a protective factor in OCCC. **a**. Kaplan-Meier curves of OCCC patients, divided by high (≥200, blue), intermediate (<200, ≥100, red) and low (<100, black) GLS1 expression levels. From left to right: PFS (12 vs 19 vs 22; *P=*0.013; HR by 10 points increase=0.93; 95% CI 0.88 to 0.99), CSS (12 vs 19 vs 21; *P=*0.014; HR by 10 points increase=0.91; 95% CI 0.85 to 0.98). OS (12 vs 19 vs 22; *P=*0.021; HR by 10 points increase =0.93, 95% CI, 0.87 to 0.99). **b**. Kaplan-Meier curves of ARID1A+ patients, divided by high (≥200, blue), intermediate (<200, ≥100, red) and low (<100, black) GLS1 expression levels. From left to right: PFS (11 vs 14 vs 10, *P=*0.023, HR by 10 points increase=0.91; 95% CI, 0.84 to 0.99), CSS (11 vs 14 vs 10, *P=*0.036, HR by 10 points increase= 0.88, 95% CI, 0.79 to 0.99), OS (11 vs 14 vs 10, *P=*0.093; HR by 10 points increase=0.93; 95% CI, 085 to 1.01). **c**. Kaplan-Meier curves of ARID1A-patients, divided by high (≥200, blue), intermediate (<200, ≥100, red) and low (<100, black) GLS1 expression levels. From left to right: PFS (1 vs 4 vs 12, *P=*0.487; HR by 10 points increase=0.96; 95% CI, 0.86 to 1.07), CSS (1 vs 4 vs 11; *P=*0.647; HR by 10 points increase=0.97; 95% CI= 0.87 to 1.09), OS (1 vs 4 vs 12; *P=*0.511; HR=0.97; 95% CI=0.87 to 1.07).

## Discussion

Clinical behavior of ovarian carcinomas is broad, ranging from highly aggressive to more indolent in nature. This behavior notably affects response to the relatively few standard-of-care treatments available for women affected by this form of cancer. Identification and validation of molecular and cellular biomarkers that can be easily tested and that are predictive of response to specific drugs remain the ideal goal of ovarian cancer research. This fact is especially crucial due to the relative dearth of drugs that induce meaningful clinical response, and even more so in the era of emerging molecular targets. Clear cell carcinomas present a particular treatment challenge. The identification of GLS1 and recent report that GLS1 is upregulated in models of ARID1A-mutant OCCC provided a basis for investigation of this target in the clinical trial setting. However, as we demonstrate here in a large tumor dataset, higher levels of GLS1 associate with better survival rates in patients with ARID1A mutations, even as ARID1A loss negatively correlated with the prognosis of these OCCC patients. GLS1 expression negatively correlated with ARID1A loss in OCCC, and its expression in ARID1A-negative OCCC is similar to its expression in normal, non-malignant ovarian and fallopian tube tissue from these patients. We detected no difference when comparing patients with early-stage vs late-stage OCCC patients.

Our findings are significant as they differ from recent reports on the association of GLS1 expression and ARID1A status. Importantly, our findings have implications for human trials using experimental therapeutics targeting GLS1. The ability to add ARID1A as a correlative predictive biomarker using IHC and/or next-generation sequencing techniques should be strongly considered for current and forthcoming trials to provide prospective validation of our findings. More importantly, in an effort to best stratify patients and limit use of targeted drugs that are not likely to be effective in patients and which may cause more harm than benefit, our findings should be examined carefully before launching iterations of this therapeutic strategy.

Limitations of this study include the retrospective approach; as always, prospective evaluation with appropriate pre-powered statistical design is indicated and necessary to confirm these findings. The number of clinical specimens used was limited to less than sixty cases. However, while a bigger cohort would be needed to validate and refine our findings, our cohort was big enough to reach significance in almost all analyses. In terms of the IHC experimental approach in the era of genomic profiling, although immunohistochemistry has been proven to be highly reliable to distinguish between *ARID1A* WT and *ARID1A*-mutated tumors^49^, the mutational status of *ARID1A* was not investigated at the RNA level. Nonetheless, this aspect does not reduce the significance of our findings given that the ARID1A protein is the one that necessarily sustains the chromatin remodeling function of ARID1A. It should be kept in mind that investigating protein levels may not be fully indicative of enzymatic activity, although an increase in GLS1 levels is always reported together with increased glutamine metabolism, including in ARID1A-mutated OCCC^19^.

## Acknowledgements

The building of the tissue microarray was funded by the Dept of Ob-Gyn & Women’s Health, UMN. This work was supported by the US Department of Defense Ovarian Cancer Research Program (OC160377), the Minnesota Ovarian Cancer Alliance, the Randy Shaver Cancer Research and Community Fund and the National Institute of General Medical Sciences (R01-GM130800) to M.B; and funding from the Minnesota Ovarian Cancer Alliance, the Randy Shaver Cancer Research and Community Fund, the Litman Family Fund for Cancer Research, the Mu Sigma Chapter of the Phi Gamma Delta Fraternity, and the American Association for Cancer Research (2019 AACR-Novocure Tumor-Treating Fields Research Grant, grant number 1-60-62-LOU) to E.L. This study was also supported by Rotary Club Forlì to V.C. We would like to thank Michael Franklin, M.S., for helpful suggestions and assistance in editing this manuscript and Dr. Jordan Mattson for the helpful discussion.

## Abbreviations

OCCC: ovarian clear cell carcinoma
ARID1A: AT-Rich Interaction Domain 1A
GLS1: glutaminase 1/ kidney isoform
SWI/SNF: SWIft/Sucrose Non-Fermentable
PFS: progression free survival
CSS: cancer specific survival
OS: overall survival

---

**Supplementary Figure 1.**
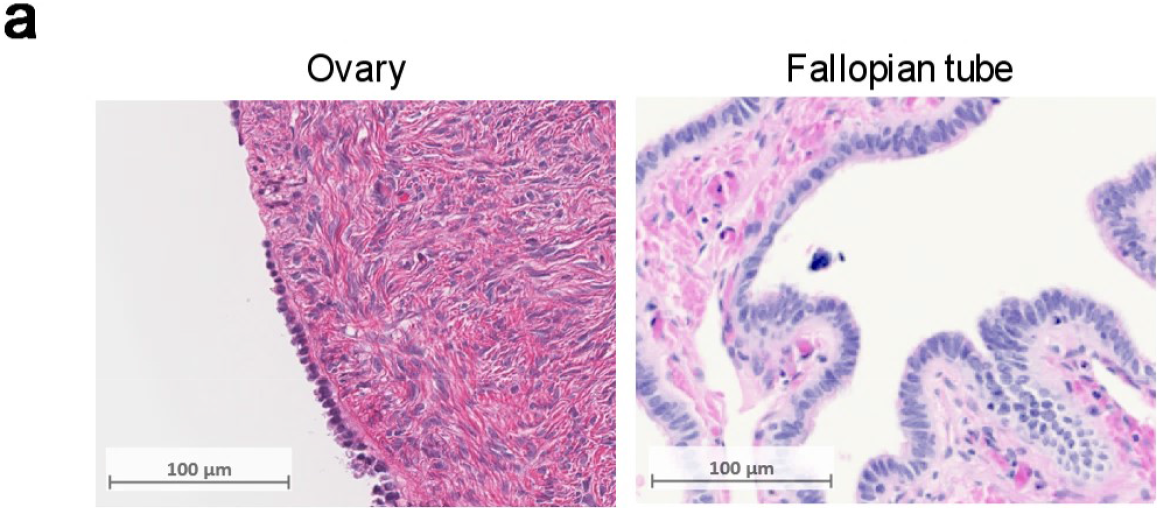
**a**. H&E staining of ovary (*left*) and fallopian tube (*right*), corresponding to the GLS1 staining in Figure 4a.

## References

1. Gadducci, A. et al. Clear cell carcinoma of the ovary: Epidemiology, pathological and biological features, treatment options and clinical outcomes. Gynecologic Oncology (2021) doi:10.1016/j.ygyno.2021.06.033.

2. Wiegand, K. C. et al. ARID1A Mutations in Endometriosis-Associated Ovarian Carcinomas. New England Journal of Medicine 363, 1532–1543 (2010).

3. Matias-Guiu, X. & Stewart, C. J. R. Endometriosis-associated ovarian neoplasia. Pathology 50, 190–204 (2018).

4. Takeda, T. et al. ARID1A gene mutation in ovarian and endometrial cancers (Review). Oncology Reports 35, 607–613 (2016).

5. Pavlidou, E. N. & Balis, V. Diagnostic significance and prognostic role of the ARID1A gene in cancer outcomes (Review). World Academy of Sciences Journal 2, 49–64 (2020).

6. Mathur, R. ARID1A loss in cancer: Towards a mechanistic understanding. Pharmacology & Therapeutics 190, 15–23 (2018).

7. Katagiri, A. et al. Loss of ARID1A expression is related to shorter progression-free survival and chemoresistance in ovarian clear cell carcinoma. Mod Pathol 25, 282–288 (2012).

8. Yokoyama, Y., Matsushita, Y., Shigeto, T., Futagami, M. & Mizunuma, H. Decreased ARID1A expression is correlated with chemoresistance in epithelial ovarian cancer. Journal of Gynecologic Oncology 25, 58–63 (2014).

9. Itamochi, H. et al. Loss of ARID1A expression is associated with poor prognosis in patients with stage I/II clear cell carcinoma of the ovary. Int J Clin Oncol 20, 967–973 (2015).

10. Jung, U. S. et al. Suppression of ARID1A associated with decreased CD8 T cells improves cell survival of ovarian clear cell carcinoma. Journal of Gynecologic Oncology 32, (2020).

11. Luchini, C. et al. Prognostic role and implications of mutation status of tumor suppressor gene ARID1A in cancer: a systematic review and meta-analysis. Oncotarget 6, 39088–39097 (2015).

12. Liu, G. et al. Prognostic and Clinicopathological Significance of ARID1A in Endometrium-Related Gynecological Cancers: A Meta-Analysis. J Cell Biochem 118, 4517–4525 (2017).

13. Luo, Q. et al. ARID1A ablation leads to multiple drug resistance in ovarian cancer via transcriptional activation of MRP2. Cancer Lett 427, 9–17 (2018).

14. Abou-Taleb, H. et al. Comprehensive assessment of the expression of the SWI/SNF complex defines two distinct prognostic subtypes of ovarian clear cell carcinoma. Oncotarget 7, 54758– 54770 (2016).

15. Ye, S. et al. Clinicopathologic Significance of HNF-1β, AIRD1A, and PIK3CA Expression in Ovarian Clear Cell Carcinoma: A Tissue Microarray Study of 130 Cases. Medicine (Baltimore) 95, e3003 (2016).

16. Zhang, X., Shetty, M., Clemente, V., Linder, S. & Bazzaro, M. Targeting Mitochondrial Metabolism in Clear Cell Carcinoma of the Ovaries. International Journal of Molecular Sciences 22, 4750 (2021).

17. Dier, U., Shin, D.-H., Hemachandra, L. P. M. P., Uusitalo, L. M. & Hempel, N. Bioenergetic analysis of ovarian cancer cell lines: profiling of histological subtypes and identification of a mitochondria-defective cell line. PLoS ONE 9, e98479 (2014).

18. Kwan, S.-Y. et al. Loss of ARID1A expression leads to sensitivity to ROS-inducing agent elesclomol in gynecologic cancer cells. Oncotarget 7, 56933–56943 (2016).

19. Wu, S. et al. Targeting glutamine dependence through GLS1 inhibition suppresses ARID1A - inactivated clear cell ovarian carcinoma. Nat Cancer 2, 189–200 (2021).

20. Yim, S. Y. et al. Low ARID1A Expression is Associated with Poor Prognosis in Hepatocellular Carcinoma. Cells 9, E2002 (2020).

21. Wang, S. C. et al. SWI/SNF component ARID1A restrains pancreatic neoplasia formation. Gut 68, 1259–1270 (2019).

22. Luo, Q. et al. ARID1A prevents squamous cell carcinoma initiation and chemoresistance by antagonizing pRb/E2F1/c-Myc-mediated cancer stemness. Cell Death & Differentiation 27, 1981–1997 (2020).

23. García-López, J. et al. Large 1p36 Deletions Affecting Arid1a Locus Facilitate Mycn-Driven Oncogenesis in Neuroblastoma. Cell Rep 30, 454-464.e5 (2020).

24. Nagl, N. G., Zweitzig, D. R., Thimmapaya, B., Beck, G. R. & Moran, E. The c-myc Gene Is a Direct Target of Mammalian SWI/SNF–Related Complexes during Differentiation-Associated Cell Cycle Arrest. Cancer Res 66, 1289–1293 (2006).

25. Shen, Y.-A. et al. Inhibition of the MYC-Regulated Glutaminase Metabolic Axis Is an Effective Synthetic Lethal Approach for Treating Chemoresistant Ovarian Cancers. Cancer Res 80, 4514–4526 (2020).

26. Szeliga, M. & Albrecht, J. Opposing roles of glutaminase isoforms in determining glioblastoma cell phenotype. Neurochem Int 88, 6–9 (2015).

27. Deng, S.-J. et al. Nutrient Stress–Dysregulated Antisense lncRNA GLS-AS Impairs GLS-Mediated Metabolism and Represses Pancreatic Cancer Progression. Cancer Res 79, 1398– 1412 (2019).

28. Zhou, W.-J. et al. Estrogen inhibits autophagy and promotes growth of endometrial cancer by promoting glutamine metabolism. Cell Commun Signal 17, 99 (2019).

29. Xiang, Y. et al. Targeted inhibition of tumor-specific glutaminase diminishes cell-autonomous tumorigenesis. J Clin Invest 125, 2293–2306 (2015).

30. Shroff, E. H. et al. MYC oncogene overexpression drives renal cell carcinoma in a mouse model through glutamine metabolism. Proc Natl Acad Sci U S A 112, 6539–6544 (2015).

31. Le, A. et al. Glucose-Independent Glutamine Metabolism via TCA Cycling for Proliferation and Survival in B Cells. Cell Metabolism 15, 110–121 (2012).

32. Craze, M. L. et al. MYC regulation of glutamine-proline regulatory axis is key in luminal B breast cancer. Br J Cancer 118, 258–265 (2018).

33. Effenberger, M. et al. Glutaminase inhibition in multiple myeloma induces apoptosis via MYC degradation. Oncotarget 8, 85858–85867 (2017).

34. Dang, C. V. MYC, microRNAs and glutamine addiction in cancers. Cell Cycle 8, 3243–3245 (2009).

35. Gao, P. et al. c-Myc suppression of miR-23a/b enhances mitochondrial glutaminase expression and glutamine metabolism. Nature 458, 762–765 (2009).

36. Dang, C. V., Le, A. & Gao, P. MYC-Induced Cancer Cell Energy Metabolism and Therapeutic Opportunities. Clin Cancer Res 15, 6479–6483 (2009).

37. Liu, W. et al. Reprogramming of proline and glutamine metabolism contributes to the proliferative and metabolic responses regulated by oncogenic transcription factor c-MYC. PNAS 109, 8983–8988 (2012).

38. Qu, X. et al. c-Myc-driven glycolysis via TXNIP suppression is dependent on glutaminase-MondoA axis in prostate cancer. Biochem Biophys Res Commun 504, 415–421 (2018).

39. Bott, A. J. et al. Oncogenic Myc Induces Expression of Glutamine Synthetase through Promoter Demethylation. Cell Metab 22, 1068–1077 (2015).

40. Chen, Z., Wang, Y., Warden, C. & Chen, S. Cross-talk between ER and HER2 regulates c-MYC-mediated glutamine metabolism in aromatase inhibitor resistant breast cancer cells. J Steroid Biochem Mol Biol 149, 118–127 (2015).

41. Xu, X. et al. Tumor suppressor NDRG2 inhibits glycolysis and glutaminolysis in colorectal cancer cells by repressing c-Myc expression. Oncotarget 6, 26161–26176 (2015).

42. Bharadwaj, S., Singh, M., Kirtipal, N. & Kang, S. G. SARS-CoV-2 and Glutamine: SARS-CoV-2 Triggered Pathogenesis via Metabolic Reprograming of Glutamine in Host Cells. Front Mol Biosci 7, 627842 (2020).

43. Thai, M. et al. MYC-induced reprogramming of glutamine catabolism supports optimal virus replication. Nat Commun 6, 8873 (2015).

44. Lukey, M. J., Wilson, K. F. & Cerione, R. A. Therapeutic strategies impacting cancer cell glutamine metabolism. Future Med Chem 5, 1685–1700 (2013).

45. Yuneva, M. O. et al. The metabolic profile of tumors depends on both the responsible genetic lesion and tissue type. Cell Metab 15, 157–170 (2012).

46. Wang, T. et al. c-Myc Overexpression Promotes Oral Cancer Cell Proliferation and Migration by Enhancing Glutaminase and Glutamine Synthetase Activity. Am J Med Sci 358, 235–242 (2019).

47. Li, J. et al. miR-145 inhibits glutamine metabolism through c-myc/GLS1 pathways in ovarian cancer cells. Cell Biology International 43, 921–930 (2019).

48. Arend, R. Phase 1 Trial of CB-839 in Combination With Niraparib in Platinum Resistant BRCA-wild-type Ovarian Cancer Patients. https://clinicaltrials.gov/ct2/show/NCT03944902 (2021).

49. Khalique, S. et al. Optimised ARID1A immunohistochemistry is an accurate predictor of ARID1A mutational status in gynaecological cancers. J Pathol Clin Res 4, 154–166 (2018).

50. Vogel, R. I. et al. USP14 is a predictor of recurrence in endometrial cancer and a molecular target for endometrial cancer treatment. Oncotarget 7, 30962–30976 (2016).

51. Shibuya, Y. et al. Identification of somatic genetic alterations in ovarian clear cell carcinoma with next generation sequencing. Genes Chromosomes Cancer 57, 51–60 (2018).

52. Lakshminarasimhan, R. et al. Down-regulation of ARID1A is sufficient to initiate neoplastic transformation along with epigenetic reprogramming in non-tumorigenic endometriotic cells. Cancer Lett 401, 11–19 (2017).

53. Chandler, R. L. et al. Coexistent ARID1A–PIK3CA mutations promote ovarian clear-cell tumorigenesis through pro-tumorigenic inflammatory cytokine signalling. Nature Communications 6, 6118 (2015).

54. Wilson, M. R., Holladay, J. & Chandler, R. L. A mouse model of endometriosis mimicking the natural spread of invasive endometrium. Hum Reprod 35, 58–69 (2020).

55. Wilson, M. R. et al. ARID1A and PI3-kinase pathway mutations in the endometrium drive epithelial transdifferentiation and collective invasion. Nature Communications 10, 3554 (2019).

56. Bulun, S. E., Wan, Y. & Matei, D. Epithelial Mutations in Endometriosis: Link to Ovarian Cancer. Endocrinology 160, 626–638 (2019).

57. Ayhan, A. et al. Loss of ARID1A expression is an early molecular event in tumor progression from ovarian endometriotic cyst to clear cell and endometrioid carcinoma. Int J Gynecol Cancer 22, 1310–1315 (2012).

